# Predictive power of non-identifiable models

**DOI:** 10.1101/2023.04.07.536025

**Authors:** Frederic Grabowski, Paweł Nałęcz-Jawecki, Tomasz Lipniacki

## Abstract

Resolving practical nonidentifiability of computational models typically requires either additional data or non-algorithmic model reduction, which frequently results in models containing parameters lacking direct interpretation. Here, instead of reducing models, we explore an alternative, Bayesian approach, and quantify predictive power of non-identifiable models. Considering an example biochemical signalling cascade model as well as its mechanical analog, we demonstrate that by measuring a single variable in response to a properly chosen stimulation protocol, the dimensionality of the parameter space is reduced, which allows for prediction of its trajectory in response to different stimulation protocols even if all model parameters remain unidentified. Successive measurements of remaining variables further constrain model parameters and enable more predictions. We analyse potential pitfalls of the proposed approach that can arise when the investigated model is oversimplified, incorrect, or when the training protocol is inadequate. The main advantage of the suggested iterative approach is that the predictive power of the model can be assessed and practically utilised at each step.

## Introduction

Non-identifiable models are ubiquitous in systems biology. Non-identifiability, in practical terms, means that parameters obtained through model fitting cannot be trusted: either because arbitrarily large changes of some parameters can be fully compensated by changes of other parameters (structural non-identifiability) or may differ widely due to noise in measurements (practical non-identifiability). It is important to note that non-identifiability is a property of a model together with a dataset, not just the model itself. Additional data can make the model identifiable^1^. Typically, structural non-identifiability is relatively easy to resolve by nondimensionalization and reparametrization. Practical nonidentifiability is a harder problem, usually requiring additional experiments (which can be expensive or time-consuming), or manual model reduction^2^. An identifiable model that reflects underlying biochemistry can be seen as the ultimate goal of systems biology. However, model reduction, performed to reach identifiability, may lead to composite parameters that lack straightforward biochemical interpretation^3–5^. Additionally, a reduced model may not be able to accommodate new data containing measurements of previously omitted variables.

In this study we focus on practically non-identifiable models, but instead of reducing them in order to reach identifiability, we propose to train them on limited datasets and investigate their predictive power after such training. Earlier Cedersund^6^ showed that non-identifiable models may lead to informative predictions, nevertheless the standard solution is to make models identifiable before predictions are made^1^, as reviewed in^2,7^. We analyse the predictive power of non-identifiable models in more detail in the context of systems biology, and investigate how successive inclusion of model variables in training influences predictions. We view modelling as the following iterative procedure: perform experiment → train model on gathered data → assess its predictive power, if required perform additional experiments → train model again, narrowing its plausible parameter space and thus increasing its predictive power. This approach stems from the fact that in (systems) biology additional measurements are usually relatively expensive and/or time-consuming, and thus researchers try to constrain the model by performing a minimal necessary set of experiments^8,9^. Ultimately, this procedure can lead to an identifiable model, however, as we will show, even without reaching identifiability, trajectories of the investigated variables can be predicted with high accuracy. To illustrate the procedure outlined above we consider a biochemical signalling cascade with a negative feedback loop—an ubiquitous motive in regulatory pathways (e.g. MAPK^10,11^, NF-κB^12,13^, p53^14,15^). As an auxiliary example we consider a simple damped mechanical oscillator consisting of three masses with one of them subject to forcing.

Model training relies on exploration of the space of plausible parameters, i.e. parameters giving predictions within an expected error bound of the measurements. There are several methods to explore the parameter space (e.g. Fisher information matrix^9,16^, profile likelihood^1,2^, Bayesian methods^17–19^, approximate Bayesian computation^20^, nested sampling^21^). We chose the Bayesian approach and sampled the parameter space performing Markov Chain Monte Carlo (MCMC) simulations using the Metropolis-Hastings algorithm^22^. This method allows us to start from broad, non-informative priors and explore the posterior parameter distributions confined by smaller or larger datasets, and consequently obtain straightforward probabilistic predictions. To avoid over-focusing on the details of the Bayesian approach, we will refer to posterior parameter samples as the set of plausible parameters, and to the process of running the MCMC simulation and obtaining the samples as training.

## Results

### Sequential training of the signalling cascade model

We consider a four step signalling cascade with negative feedback(s) resembling the MAPK cascade (RAS → RAF → MEK → ERK), outlined in Figure 1a, with corresponding equations given in Figure 1b. The cascade is activated by a time dependent signal *S*(*t*), which depends on the stimulation protocol chosen. Nominal model parameters (used for generating training data) and assumed lognormal parameter priors used during model training are given in Figure 1c. We note that, by default, the strengths of feedbacks to the second and third step of the cascade are zero (*f*_2_ = *f*_3_ = 0)—later we investigate a ‘relaxed’ model in which *f*_2_ and *f*_3_ are allowed to be positive. The training dataset is generated by simulating trajectories of the four model variables, *K*_1_, *K*_2_, *K*_3_, *K*_4_ using an ‘on-off’ stimulation protocol and randomly perturbing them in order to mimic experimental errors (Fig. 1d). The error bars are plotted at measurement time points (every hour) and show the magnitude of the assumed experimental errors. We use three such generated measurement replicates for training (see Methods for details). The training dataset is used to train two models: the ‘nominal’ model for which we assume *f*_2_ = *f*_3_ = 0, and a ‘relaxed’ model (in the next section) in which we allow *f*_2_ and *f*_3_ to assume positive values, also with the lognormal prior (Fig. 1c).

**Figure 1.**
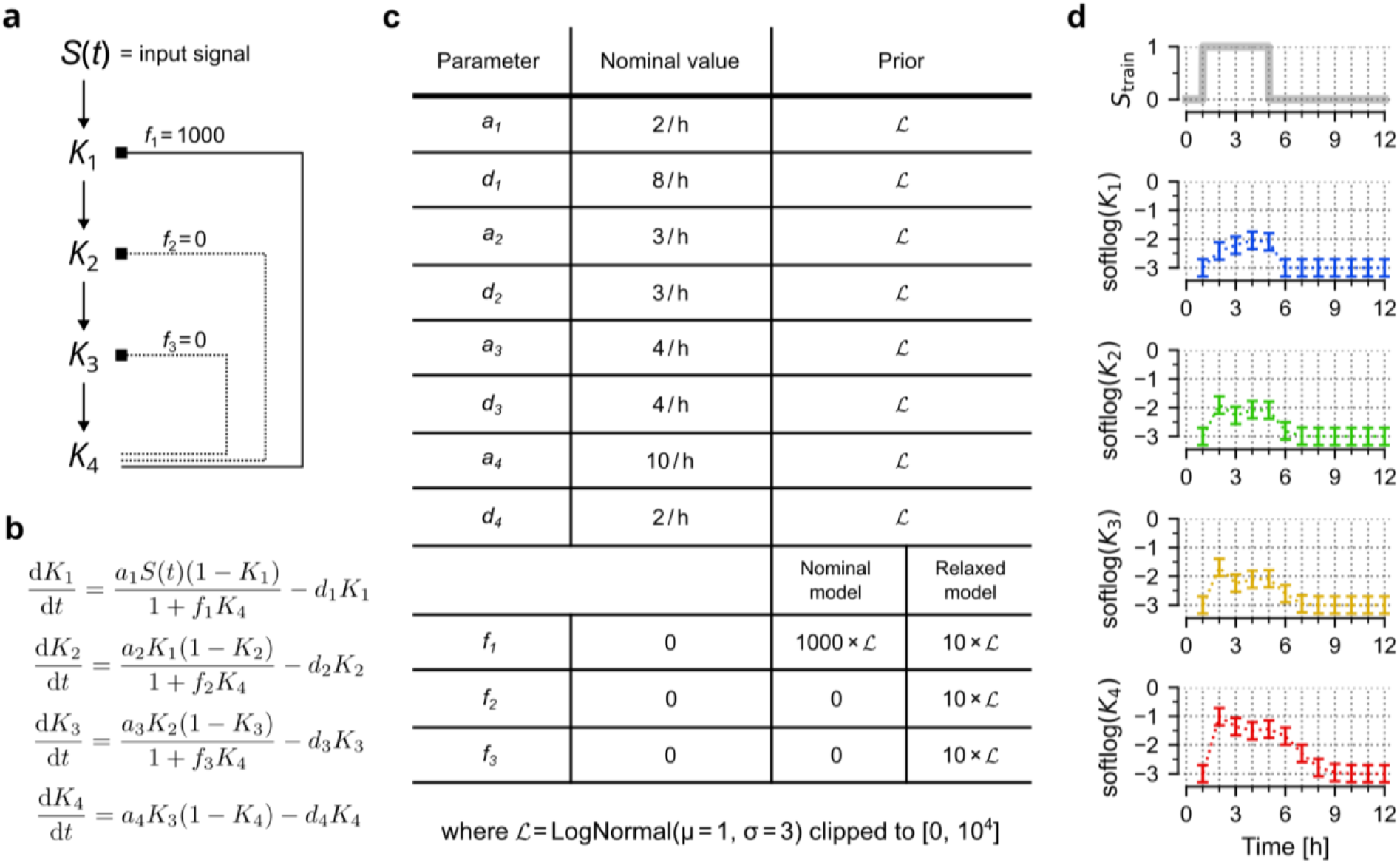
Signalling cascade model. (**a**) Scheme of the 4-level biochemical signalling cascade model activated by a time-dependent signal *S*(*t*). Strengths of feedbacks *f*_2_ and *f*_3_ are set to zero in the nominal model, but are allowed to assume positive values in the ‘relaxed’ model. (**b**) Model equations. (**c**) Model parameters used for generating training data and prior distributions used for training. (**d**) Simulated trajectories of *K*_1_, *K*_2_, *K*_3_, *K*_4_ in response to *S*_train_ used for model training. Error bars show measurement errors assumed for generation of training data.

First, we train the nominal model using only the trajectory of the last variable, *K*_4_. We observe that the resulting model accurately predicts the trajectory of *K*_4_ in response to two very different stimulation protocols (Fig. 2a and 2b), but fails to predict trajectories of any of the remaining variables. For these three variables the 80% prediction bands are very broad, implying that the trained model contains little information about these variables. Next, the training dataset is expanded to additionally include the trajectory of *K*_2_, and we observe that also the trajectory of this variable can be reliably predicted (Fig. 2c). Finally, the complete training dataset containing trajectories of all four variables results in a “well trained” model, which predicts the trajectories of all variables (Fig. 2d).

**Figure 2.**
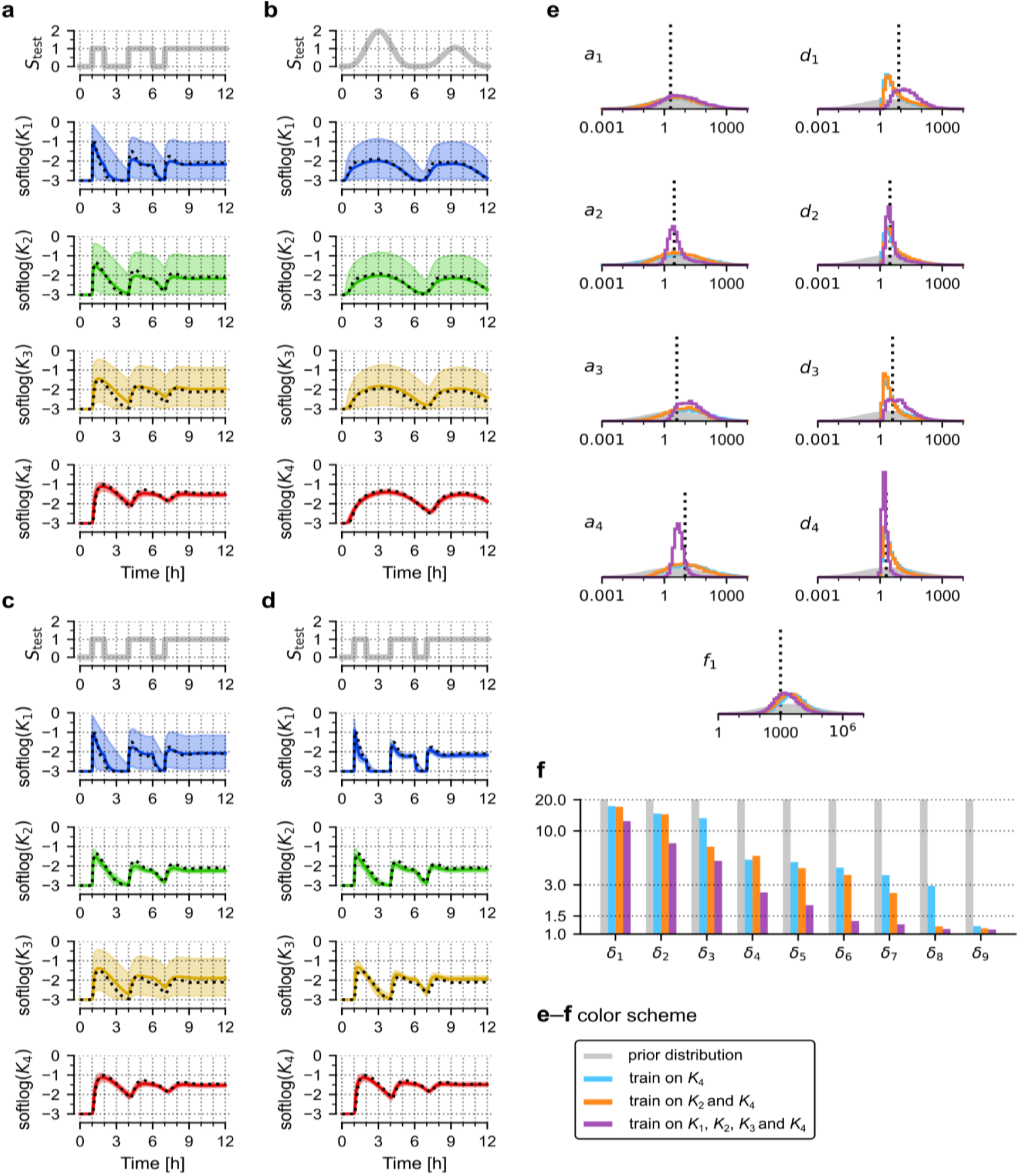
Training the nominal model. (**a**–**b**) Prediction of model responses to two test signals, *S*_test_, after training on trajectory of *K*_4_ (generated in response to the training protocol, *S*_train_, shown in Figure 1d). Black dotted lines show trajectories of the nominal model, coloured lines show (point-wise) medians of predictions, contours show 80% prediction bands. (**c**) Prediction of model responses after training on trajectories *K*_2_ and *K*_4_. (**d**) Prediction of model responses after training on trajectories of all four model variables, *K*_1_, *K*_2_, *K*_3_ and *K*_4_. (**e**) Histograms obtained from 10,000 samples generated from the posterior distribution of model parameters after training on *K*_4_ (blue); *K*_2_ and *K*_4_ (orange); *K*_1_, *K*_2_, *K*_3_ and *K*_4_ (purple). Grey contours show prior distributions. (**f**) Dimensionality reduction of the parameter space by successive inclusion of model variable trajectories into the training set. Bar heights show principal multiplicative deviations δ_i_ = exp(√λ_i_), where λ_i_ are principal values (variances) of log-parameters.

By investigating the space of plausible parameters (i.e., posterior samples) we may notice that even for the well trained model the marginal probability distributions show that all 9 parameters may vary by about an order of magnitude, and when training is performed only on *K*_4_ (or *K*_2_ and *K*_4_) parameters may vary by about two orders of magnitude (Fig. 2e). This shows that the model trained only on the last variable of the cascade, *K*_4_, can accurately predict this variable’s trajectories in response to different stimulation protocols, even though all model parameters remain uncertain. To better understand this feature we numerically analyse the dimensionality of the plausible parameter sets (see Methods for details) obtained after training the model on *K*_4_, *K*_2_ and *K*_4_, and all four variables (Fig. 2f). Since we are interested in multiplicative changes of parameters, we perform principal component analysis (PCA) on sets of (natural) logarithms of plausible parameters. Exponents of the square roots of the principal values (variances) λ_i_ can be interpreted as principal multiplicative deviations of parameter values, δ_i_ = exp(√λ_i_). The prior parameter distributions are LogNormal(μ = 1., σ = 3.) clipped to [0, 10^4^], which gives δ_i,prior_ ≈ exp(3.) ≈ 20 (for all *i*), implying that “on average” all prior parameter values can vary 20-fold from the median. After training on all variables the largest principal multiplicative deviation is δ_i_ ≈ 12, however, the 4 smallest δ_i_ are much lower, below 1.5 (note that δ_i_ = 1 implies no variation along the *i*-th principal component). This implies that the dimensionality of the plausible parameter space was effectively reduced by 4 due to training. When training is performed only on *K*_4_, dimensionality is reduced by 1 (only the smallest δ_i_ is below 1.5), and when training uses *K*_2_ and *K*_4_, dimensionality is reduced by 2 (two smallest δ_i_ are below 1.5). Therefore, in agreement with intuition, model training reduces dimensionality of the plausible parameter space. In our example, the measurement of *n* variables reduces dimensionality by *n*. Importantly, even reduction from 9 to 8 dimensions allows for reliable predictions of a single variable of interest that was measured to train the model.

We performed a similar analysis of a (distant) mechanical analogue of the signalling pathway, a damped harmonic oscillator consisting of three masses connected by two springs (Fig. S1). The system is parametrized by 8 free parameters. For training we use a protocol in which one of the masses is pushed with constant force for 10 seconds, which results in a displacement of all masses. Next, we used the trajectory of the opposite mass (or of all masses) to train the model. In the test protocol, a sinusoidal driving force was applied to the same mass as during training. Similarly to the signalling cascade model, we found that training on a single mass trajectory suffices to predict its trajectory (for a limited time) in response to a very different forcing, and training on all three masses allows for very accurate predictions of all trajectories, despite the fact that marginal distributions of model parameters remain broad.

### Training a relaxed model

Frequently, experiments are performed not only to constrain parameters of a known model, but to deduce the structure of the model itself. To explore this scenario, we introduce a ‘relaxed’ model, in which we allow strengths of the three negative feedbacks shown in Figure 1a to assume positive values during training (assuming prior lognormal distributions, as shown in Figure 1c). This increases the number of free parameters from 9 to 11. Again, this relaxed model is trained on *in silico*-generated data, based on the nominal model with one feedback. Regardless of whether only *K*_4_ or all variables are included in the training dataset, the relaxed model performs equally well as the nominal model on test protocols (compare Fig. 3A versus Fig. 2A and Fig. 3B versus Fig. 2D). We note that the marginal distributions of the three feedback parameters remain broad (Fig. 3c). This means that, even without knowing strengths of individual feedbacks, the model is able to give reasonable predictions. Nevertheless, these distributions indicate that the (nominal) feedback *f*_1_ is dominant.

**Figure 3.**
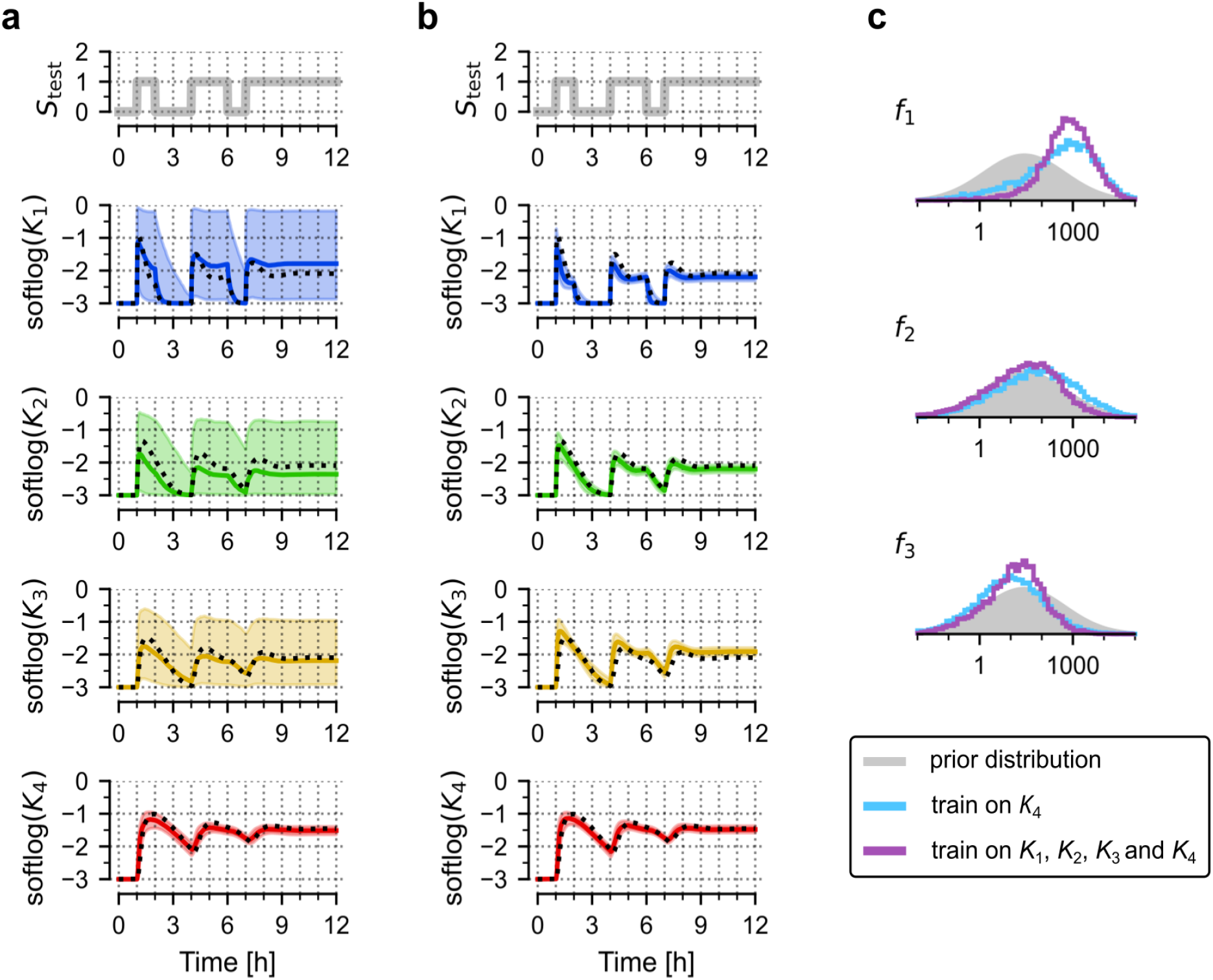
Training a relaxed model. (**a**) Prediction of model responses to test signal, *S*_test_, after training on trajectory of *K*_4_ (generated in response to the training protocol, *S*_train_, shown in Figure 1d). Black dotted lines show trajectories of the nominal model, coloured lines show (point-wise) medians of predictions, contours show 80% prediction bands. (**b**) Prediction of model responses after training on trajectories of all four model variables. (**c**) Histograms obtained from 10,000 samples generated from the posterior distribution of model parameters after training on *K*_4_ (blue) or all model variables (purple). Grey contours show prior distributions.

The above example suggests that if there exists uncertainty about model structure, it is advisable to train a more relaxed model, which can still result in satisfactory predictions, and eventually, after gathering more data, can provide a hint about true model structure.

### Training a simplified model

Given insufficient knowledge about the cascade structure and a small dataset, it might be tempting to train a simplified model. Here, we give an example that showcases potential pitfalls of this approach. As previously, the training set is a perturbed trajectory of *K*_4_ generated by the nominal model—however, we will attempt to train a model that contains only two signalling steps, in which *K*_1_ directly influences *K*_4_ (formally in equation for *K*_4_ we replace *K*_3_ by *K*_1_, and equations for *K*_2_ and *K*_3_ are omitted). Training the simplified model on the ‘on-off’ protocol shows some discrepancy between original and simplified model trajectories (Fig. 4a, note the logarithmic scale of the plot). When the trained model is verified on the test stimulation protocol, the observed discrepancy is more visible, meaning that the simplified model is not able to make accurate predictions (Fig. 4b). Crucially, the 80% prediction band is relatively narrow, similar to that obtained for the correct model (Fig. 4b versus Fig. 2a). Thus the trained simplified model is “sure” (even if marginal parameter distributions of 3 out of 5 model parameters remain very broad—Fig. 4c) about its inaccurate predictions.

**Figure 4.**
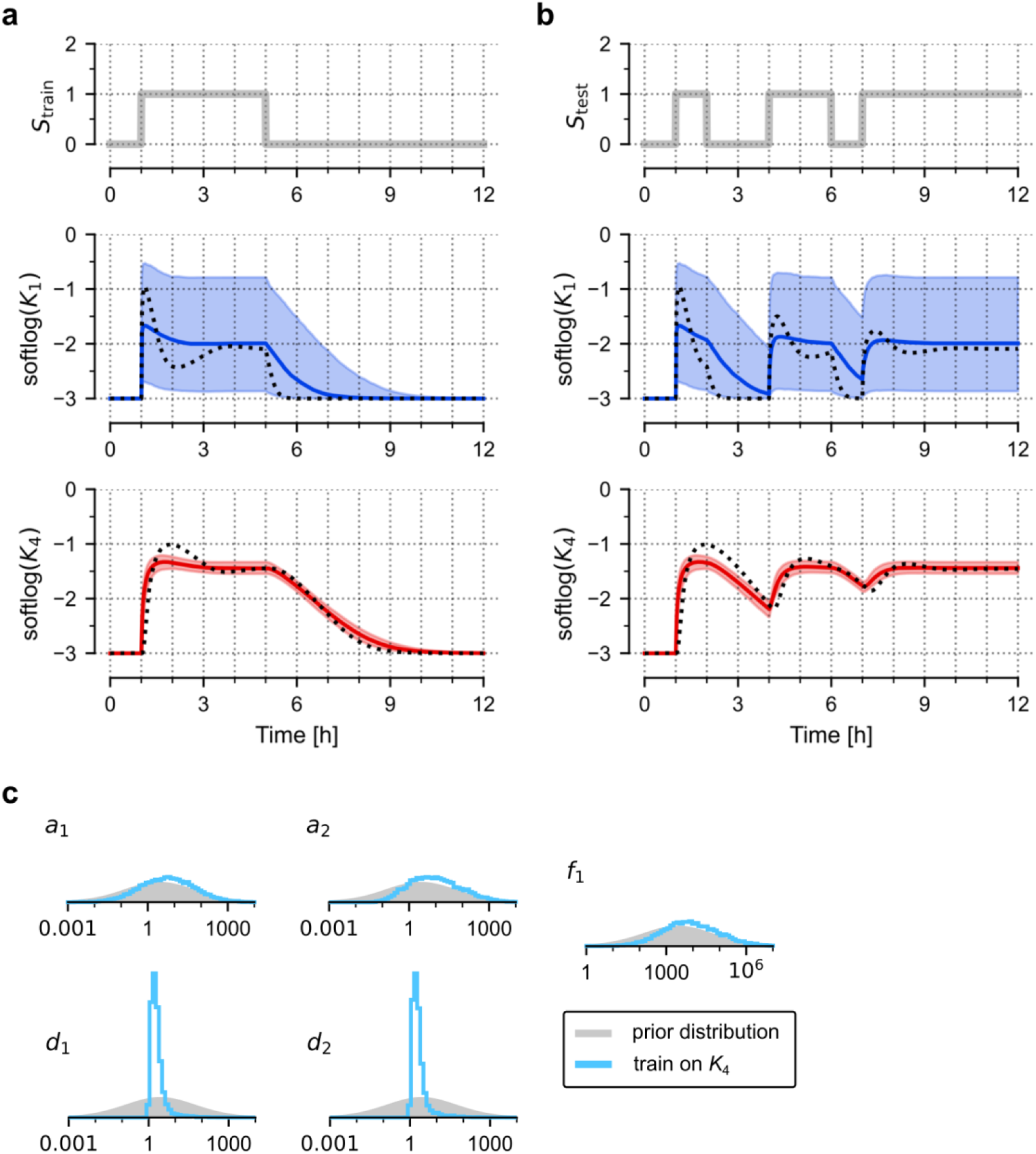
Training a simplified model. (**a**) Training a simplified (2-step) model on the nominal (4-step) model trajectory of *K*_4_. Only variables present in the simplified model are shown. Black dotted lines show trajectories of the nominal model, coloured lines show (point-wise) medians of the simplified model, contours show 80% prediction bands. (**b**) Prediction of the trained simplified model responses to the test signal, *S*_test_. Same graphical convention as in (**a**). (**c**) Histograms obtained from 10,000 samples generated from the posterior distribution of simplified model parameters. Grey contours show prior distributions.

In certain cases where approximate predictions are still satisfactory, the simplified model can be a reasonable solution. However, as demonstrated above, one should be very careful when interpreting parameters and making predictions, since some conclusions might be an artefact of using a wrong model to compute posterior probabilities. Finally, in contrast to the relaxed model, the simplified model cannot be improved by additional data.

### Analysis of potential pitfalls

In Figure S2 we investigate training based on an inadequate stimulation protocol. The protocol’s ‘on’ phase lasts only 1 hour (compared to 4 hours in Figure 2), and as a result the model predictions have a broad 80% prediction band. The prediction band for *K*_4_ is broad when training includes *K*_4_ only, and remains broad even when training includes all four model variables. Higher uncertainty of trained model trajectories is associated with larger principal multiplicative deviations δ_i_, compared to values obtained for training shown in Figure 1d. This example highlights the importance of designing a good stimulation protocol for training. If our goal is to obtain accurate predictions of *K*_4_ only, training on *K*_4_ with help of an appropriate protocol (Fig. 2a) gives much better results than training on all variables employing an inadequate protocol (Fig. S2c). This pitfall is relatively simple to resolve—one has to adjust the training protocol, so that it allows capturing characteristic temporal responses of the system. Whether the 1 hour long ‘on’ phase is sufficient or not depends on the nominal model parameters.

In Figure S3 we show an example where the prior model is incorrect, i.e. has no negative feedback at all. We obtain discrepancies between measurement and model, which could be overlooked during training if, for example, the measurement points were not distributed densely enough. From the trained wrong model we also obtain wrong predictions with very narrow prediction bands, as in the case of the simplified model discussed earlier. The key message from Figure 4 and Figure S3 is that a simplified or wrong model can make inaccurate or wrong predictions with narrow prediction bands, and thus the width of prediction bands contains no information about accuracy of the model. Narrow prediction bands imply that obtained constraints on parameters suffice for unique predictions, but these predictions are correct only under the condition that the model structure is also correct.

## Discussion

Majority of systems biology models, including mechanistic models of intensely studied regulatory networks, are non-identifiable. Developed techniques allow to automatically resolve structural non-identifiability, however, resolving practical non-identifiability is harder, and frequently requires additional data or substantial model reduction. We see the model reduction problem as follows: regulatory networks have built-in signalling cascades that transmit signals between key nodes. In many cases researchers are aware of components (typically kinases) of these signalling cascades, but measuring kinetics of all these components is not feasible. It is thus tempting to skip these intermediate steps, but this solution is problematic since these intermediate steps introduce time delays that can be important for whole-network behaviour, such as oscillations. In the case of the discussed cascade *S → K*_1_ *→ K*_2_ *→ K*_3_ *→ K*_4_ triggered in *t* = 0 from state *K*_1_ *= K*_2_ *= K*_3_ *= K*_4_ = 0 by the step signal *S*(*t*), the first, second and third time derivatives of *K*_4_ (at *t* = 0) are zero. When the cascade is reduced to *S → K*_1_ *→ K*_4_, the second derivative of *K*_4_ becomes positive, which noticeably changes the system’s response, as shown in Figure 4. To maintain observed dynamics (in particular oscillations) one can introduce nonlinearities, but as a consequence the resulting model misses some components and has nonlinear terms not supported by underlying biochemistry^23,24^.

In response to the outlined problems with model reduction, we proposed an alternative approach based on training the full mechanistic model on the available dataset and assessing its predictive power. We have demonstrated that a very limited dataset, insufficient to constrain any model parameters, can still allow for satisfactory predictions. In particular, we showed that by training the signalling cascade model on the trajectory of the last component of the cascade (obtained in response to some properly chosen training protocol), one can unambiguously predict its trajectory in response to other protocols. This prediction is possible despite the fact that trajectories of all upstream variables remain unknown. In the considered example we observe that when training is performed on trajectories of *n* variables, the ‘effective’ dimensionality of the plausible parameter set is reduced by *n*, and this reduction suffices to predict trajectories of the measured variables in response to other protocols. As discussed by Matcha et al. compression of the parameter space identifies sensitive or stiff directions corresponding to the composite parameters controlling observed aspects of system behaviour^25^.

compression of the microscopic parameter space, with sensitive or stiff directions corresponding to the relevant macroscopic parameters (such as the diffusion constant). These results suggest that the hierarchy of theories in physics relies on the same parameter space compression that is ubiquitous in general multiparameter models.

Model training can be generally understood as exploring the parameter space, to find plausible parameters, i.e these leading to model trajectories in agreement with available data. In this context, many exploration methods are available. Locally, information about uncertainty of parameters can be calculated using Fisher’s information matrix. For models with asymmetric but still unimodal loss landscapes the (semi-local) profile likelihood method can be used. For arbitrary loss landscapes, a Bayesian approach together with MCMC exploration is convenient, as

1. it allows for inclusion of prior knowledge; if no such knowledge exists, training can start from some broad uninformative prior,
2. it allows to analyse how much a model has learnt by including a given dataset; this can be expressed by the reduction of the ‘effective’ dimensionality of the plausible parameter space, or entropy of plausible parameters distribution,
3. It allows us to assess the predictive power of the trained model, by sampling the posterior parameter distribution and computing prediction bands for variable trajectories

The main disadvantage of MCMC compared to local (or semi-local) methods is that it can become computationally expensive, especially for high-dimensional parameter spaces^19^. The computational cost is typically lower if exploration starts from a relatively narrow (informative) prior, or a more complete dataset is available, which results in a narrower posterior distribution^26^. A promising method of accelerating MCMC exploration is based on reducing expensive model evaluations by local approximation of the likelihood^27,28^.

In summary, the advantage of the proposed approach to non-identifiability is twofold.

1. It does not require to preemptively reduce models to avoid unmeasured variables in order to obtain predictions. Certainty of the predictions obtained from the full model can be determined by predictive bands, and if these predictions are insufficiently narrow, the model can be further trained using additional measurements of the variables of interest. Notably, these predictions are more reliable than predictions obtained from simplified models. While predictions from the full model may have wide predictive bands, the predictions from simplified models can be misleading (inaccurate with narrow bands) due to introduced simplifications.
2. If at a later time measurements of previously unmeasured variables are performed, no modification of the model structure is necessary. In contrast, reduced models may not be able to accommodate new data, simply because some components were omitted during reduction.

We hope that this approach will allow for easier collaboration, as it allows researchers to jointly work on the same model, improving it by performing new experiments.

Identifiable models or theories stem from physics, where they showed their superiority over other approaches. Whole branches of physics are based on a small number of fundamental theories. In systems biology, the situation is different: there are a lot of not-so-important models, typically having a large number of parameters that can neither be (feasibly) derived from more fundamental theories nor effectively measured. Currently, a very intensively developed approach to complex systems is based on neural networks. In this approach one gets little or no insight into how the system works, but can get very good predictions. In this sense, Bayesian methods are a middle ground between classical, identifiable models and black box models such as neural networks. The proposed Bayesian approach works well in our case because we have a good prior on the model structure, even if the prior on model parameters is broad and uninformative. This approach can be impractical for very complex systems for which neural networks are typically used, but is able to both give useful predictions and obtain some insight into the underlying regulatory system. Even if the explanatory power of the trained non-identifiable model is limited by uncertainty of its parameters, its predictive power can make it useful (see discussion in^29^).

We think that the proposed approach, based on Bayesian inference from an incomplete dataset that gives partial insight into the system’s behaviour, may reflect true uncertainties in underlying biology. Considering a regulatory network one can identify core nodes. Because of their importance, the corresponding variables are frequently measured, and for the same reason one can expect relatively good replicability of these measurements. The intermediate pathway components are less tightly constrained, and may exhibit higher variability. This implies that (for very practical reasons) these variables are not (frequently) measured; or even if they are measured, results are not reported due to insufficient replicability. In other words, in the case of intensively investigated regulatory systems, uncertainty of model parameters and of trajectories of intermediate components may reflect their true variability, at least at the single cell level.

## Methods

### Computational model simulations

Each model is represented by a set of ODEs and an input signal *S*(*t*), that are solved using an explicit Runge-Kutta (4,5) formula developed by Dormand-Prince. Model simulations are used first to generate *in silico* measurements, then to estimate parameters distributions based on these measurements. A measurement trajectory is evaluated at full hours (or at full seconds in the case of the mechanistic analog model shown in Fig. S1) by adding to the ODE solution gaussian noise (with sigma σ_error_ = 0.3) that emulates measurement error. Parameter estimation (model training) is based on 3 independent measurement trajectories.

In the case of biochemical cascade models, ODE trajectory *x*(*t*) is log-transformed to *y*(*t*) = log_10_(0.001 + *x*) before adding measurement noise.

### Parameter estimation based on MCMC simulations

For each of the considered models we assume some prior parameter distribution *P*(θ) and use an MCMC algorithm to draw samples from the posterior parameter distribution *P*(θ | *M*), given *in silico* generated measurements *M*.

To estimate *P*(θ | *M*), we first calculate the likelihood of θ given a single measurement *L*_k,t,r_(θ) = *P*(*m*_k,t,r_ | θ) (where *k* is the measured variable, *t* is the time of measurement, and *r* = 1, 2, 3 is the measurement replicate), we simulate variable trajectories *y*_k_(*t*) for a parameter set θ. Then

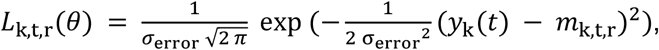

where σ_error_ is the measurement error (assumed 0.3 for all our models). The total likelihood given all measurements of variable *k* is thus:

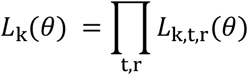

and the posterior distribution *P*(θ | *M*) is proportional to

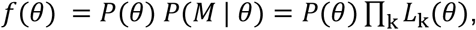

where the product is over variables k selected for model training. Then *f*(θ) is used in MCMC simulations to draw samples from *P*(θ | *M*).

MCMC simulations are performed using the Metropolis-Hastings algorithm with Gaussian jumping distribution. We use 8 parallel chains, each with 1.5 million steps from which the first 0.25 million are removed (burn-in). This gives altogether 10 million parameter samples from which we retain every thousandth sample, i.e. 10,000 samples for model simulations. We compared the between- and within-chain variance estimates using the r-hat convergence statistics to find r-hat < 1.01, suggesting sufficient length of chains. The effective sample size of the final sample was found to be at least 1,000.

For the biochemical cascade models, we assume wide log-normal priors and thus use log-parameters during MCMC sampling. After this reparametrization, we use a spherically symmetric Gaussian jumping distribution with independent variables and σ = 0.2 (in the log scale). For the mechanical model we use an elliptical Gaussian jumping distribution with σ = 1.0 for masses, σ = 0.2 for damping coefficients and σ = 0.4 for stiffness coefficients. For the final fit (trajectories of all three masses observed) we decrease step size by a factor of 4.

To minimise biases we select wide, non-informative priors, however it is worth noting that in practical applications informative priors can significantly increase model performance.

In summary, our statistical model can be itemised as follows:

1. *θ*∼ Prior distribution
2. *x*(*t*) = Model ODEs solution for parameters *θ* and initial condition *x*(0) = *x*_0_
3. *y*(*t*) = *softlog*(*x*(*t*)) := (0.001 + *x*(*t*)) (*only for signalling cascade)*
4. *m*_k,t,r_ ∼ 𝒩(*y*_k_(*t*), σ_error_)

where θ are the model parameters, σ_error_ is the measurement error and *m*_k,t,r_(*t*) is the *r*-th measurement of variable *k* at time *t*, drawn from normal distribution 𝒩(*y*_k_(*t*), σ_error_). Initial conditions for the ODE are kept constant and excluded from estimation. The *softlog* normalisation step is omitted for spring models. For all our models σ_error_ = 0.3.

### Principal component analysis

In Figure 3C and Figure S2D we use principal component analysis (PCA) to quantitatively describe dimensionality reduction of the plausible parameter set. This approach works if the analysed set of points is “sufficiently flat”. Points on a half circle would have two big PC values, even though the space is clearly one dimensional. In such a case dimensionality can be investigated locally, as proposed by Little et al.^30^. In the half circle example, this local PCA analysis would result in one small and one big PC value (for a sufficiently small spatial scale). We have verified that in our case local estimates result in higher PC values than the global ones; we thus conclude that our point clouds are sufficiently flat to justify using global PCA analysis.

### Implementation

We implemented our models in Python using jax^31^ and diffrax^32^ (for ODE integration), and the MCMC algorithm using blackjax^33^.

## Code availability

Our source code is available at https://github.com/grfrederic/identifiability.

## Acknowledgements

This study was funded by National Science Centre Poland grants 2019/34/H/NZ6/00699 (Norwegian Financial Mechanism) and 2019/35/B/NZ2/03898. We thank Dr. Marek Kochanczyk for discussions.

## Author contributions

F.G., P.N-J. and T.L. designed the study, F.G. wrote numerical codes, performed simulations and prepared figures. F.G. and T.L with P.N-J. assistance wrote the manuscript.

## Data availability

No experimental datasets were generated or analysed during the current study.

All numerical results were obtained using code available at https://github.com/grfrederic/identifiability.

## Additional Information

### Supplementary information

Includes supplementary figures S1–S3.

### Appendix

**Figure S1.**
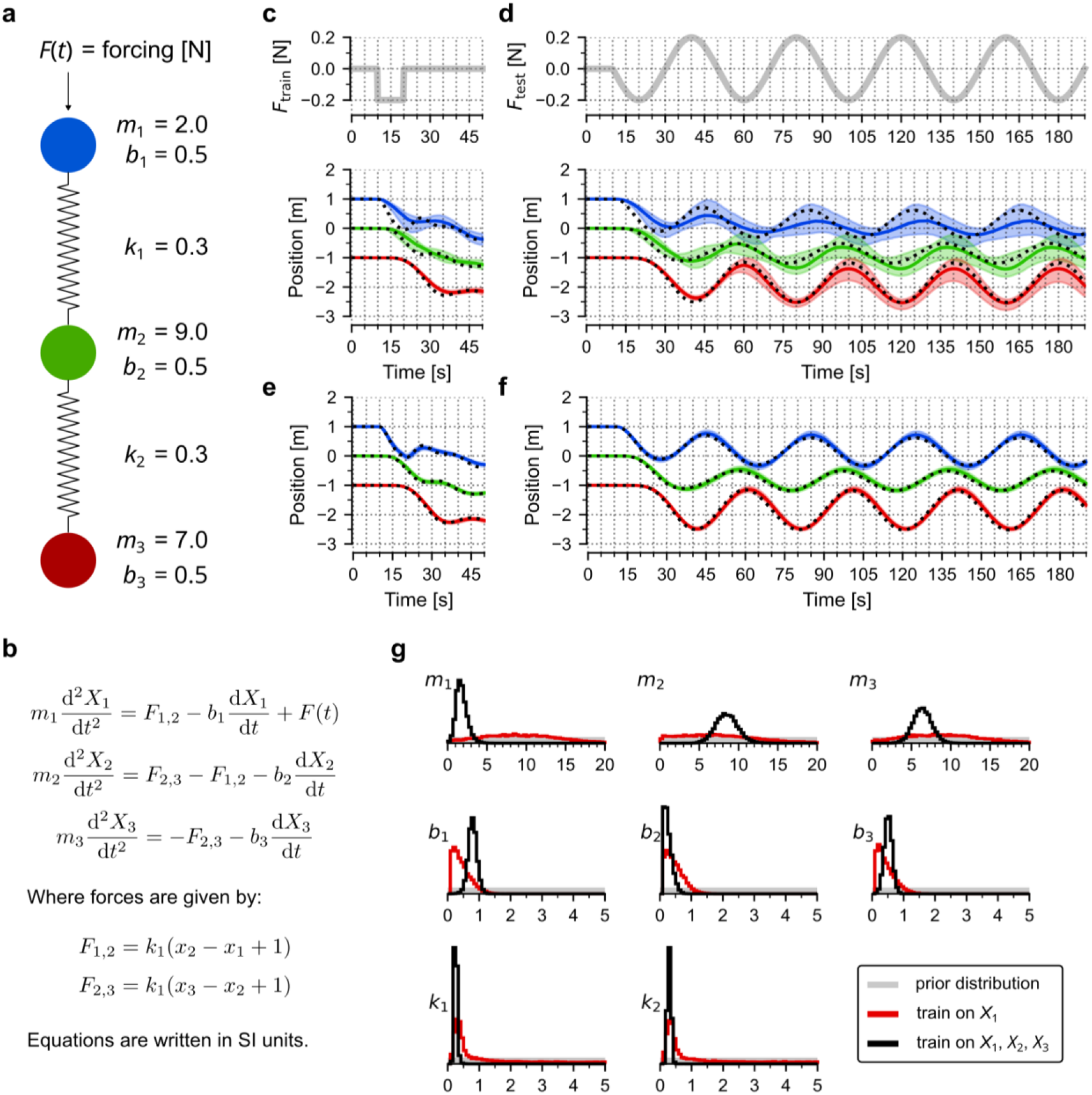
Three-body harmonic oscillator. (**a**) Scheme of the model. All eight parameters (masses, *m*_i_; damping coefficients *b*_i_; spring constants *k*_i_) to be estimated are given together with their nominal values. Before the time dependent force *F*(*t*) is applied, the system remains in equilibrium. (**b**) Model equations. (**c**–**d**) Training (**c**) and prediction (**d**), with only the trajectory of the red mass included in training. As in the signalling cascade model, for training we use three independently generated measurement replicates with gaussian noise (sigma σ_error_ = 0.3) added to the deterministic trajectory (black dotted line). Measurement points are at full seconds. Coloured lines show (point-wise) medians of predictions, contours show 80% prediction bands. (**e**–**f**) Training (**e**) and prediction (**f**), with trajectories of all masses included in training, same convention as in **c**–**d**. (**g**) Histograms obtained from 10,000 samples generated from the posterior distribution of model parameters after training on red mass trajectory (red line) and after training on all masses trajectories (black line). Grey contours show (uniform) prior distributions.

**Figure S2.**
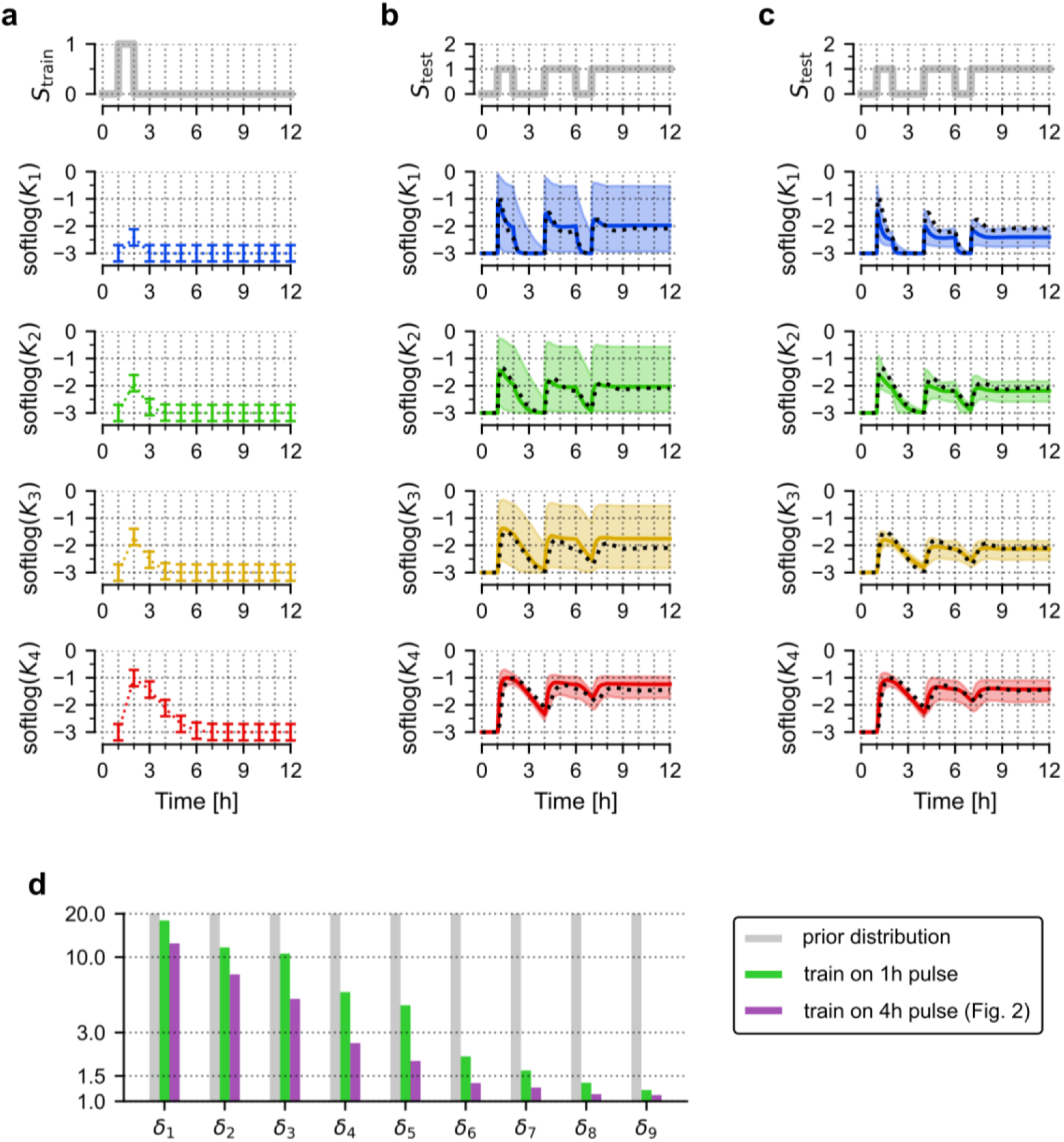
Training with an inadequate protocol. (**a**) Simulated trajectories of *K*_1_, *K*_2_, *K*_3_, *K*_4_ in response to a shorter pulse (1h instead of 4h) *S*_train_ used for model training. Error bars show measurement errors assumed for generation of training data. (**b**–**c**) Prediction of model responses to a test signal *S*_test_ after training on trajectory of *K*_4_ (**b**); and all four model variables (**c**). Black dotted lines show trajectories of the nominal model, coloured lines show (point-wise) medians of predictions, contours show 80% prediction bands. Including additional variables in training does not increase the model’s predictive power for *K*_4_. (**d**) Dimensionality reduction of the parameter space by training on all four model variables and different pulse lengths. Purple—training on a 4h pulse (repeated from Figure 2), green—training on a 1h pulse, grey— prior. Compared to training on a 4h pulse, principal multiplicative deviations are larger.

**Figure S3.**
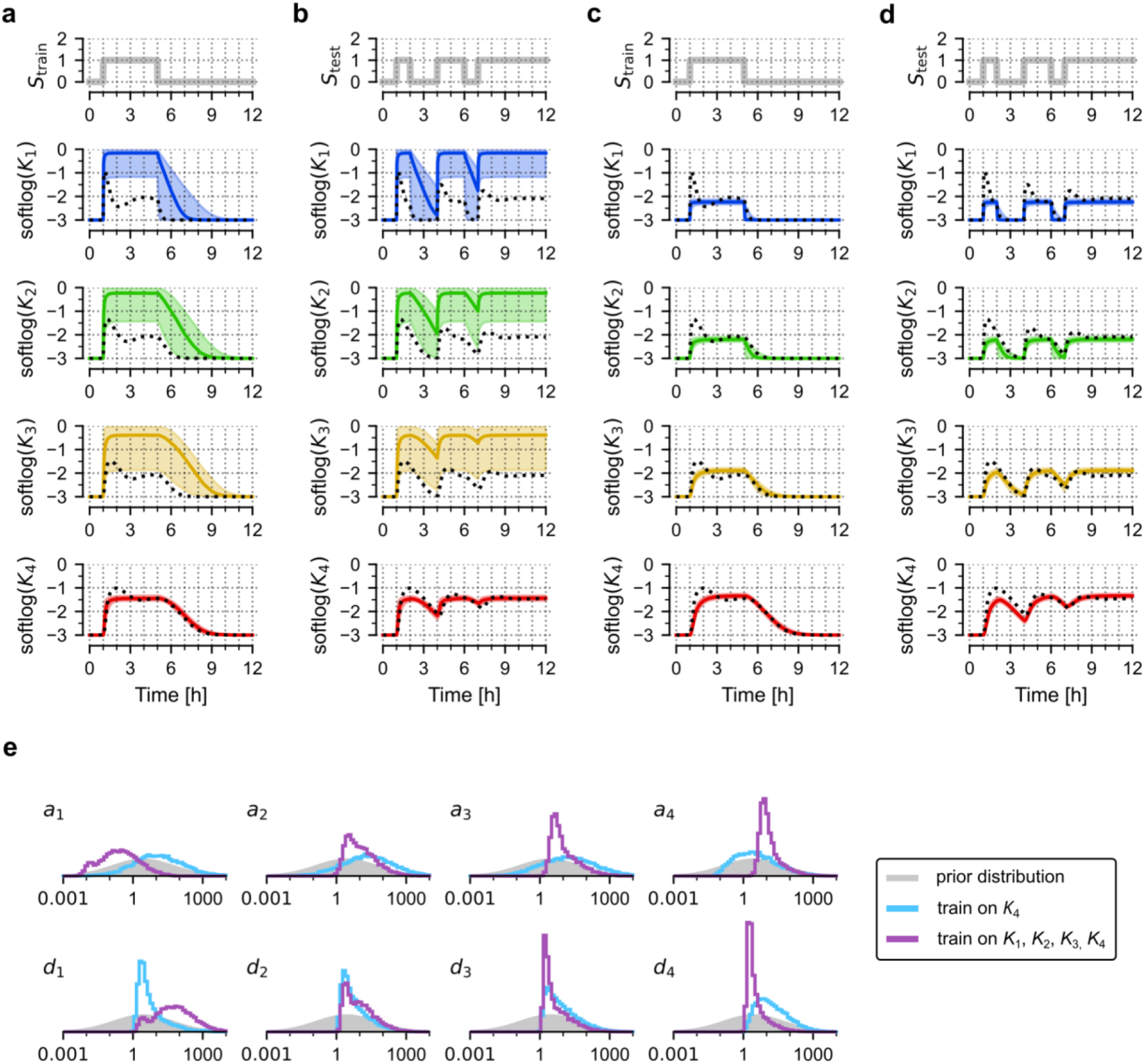
Training an incorrect model (without feedback). (**a**–**d**) Prediction of model responses to the train signal *S*_train_ (**a, c**) and a test signal *S*_test_ (**b, d**) after training on trajectory of *K*_4_ (**a, b**) and all four model variables (**c, d**). Black dotted lines show trajectories of the nominal model, coloured lines show (point-wise) medians of predictions, contours show 80% prediction bands. Model confident and wrong. (**e**) Histograms obtained from 10,000 samples generated from the posterior distribution of the incorrect model parameters after training on *K*_4_ (blue); *K*_1_, *K*_2_, *K*_3_ and *K*_4_ (purple). Grey contours show prior distributions.

